## Supplementary material for "Systematic Allelic Analysis Defines the Interplay of Key Pathways in X Chromosome Inactivation"

#### Supplementary Figure 1

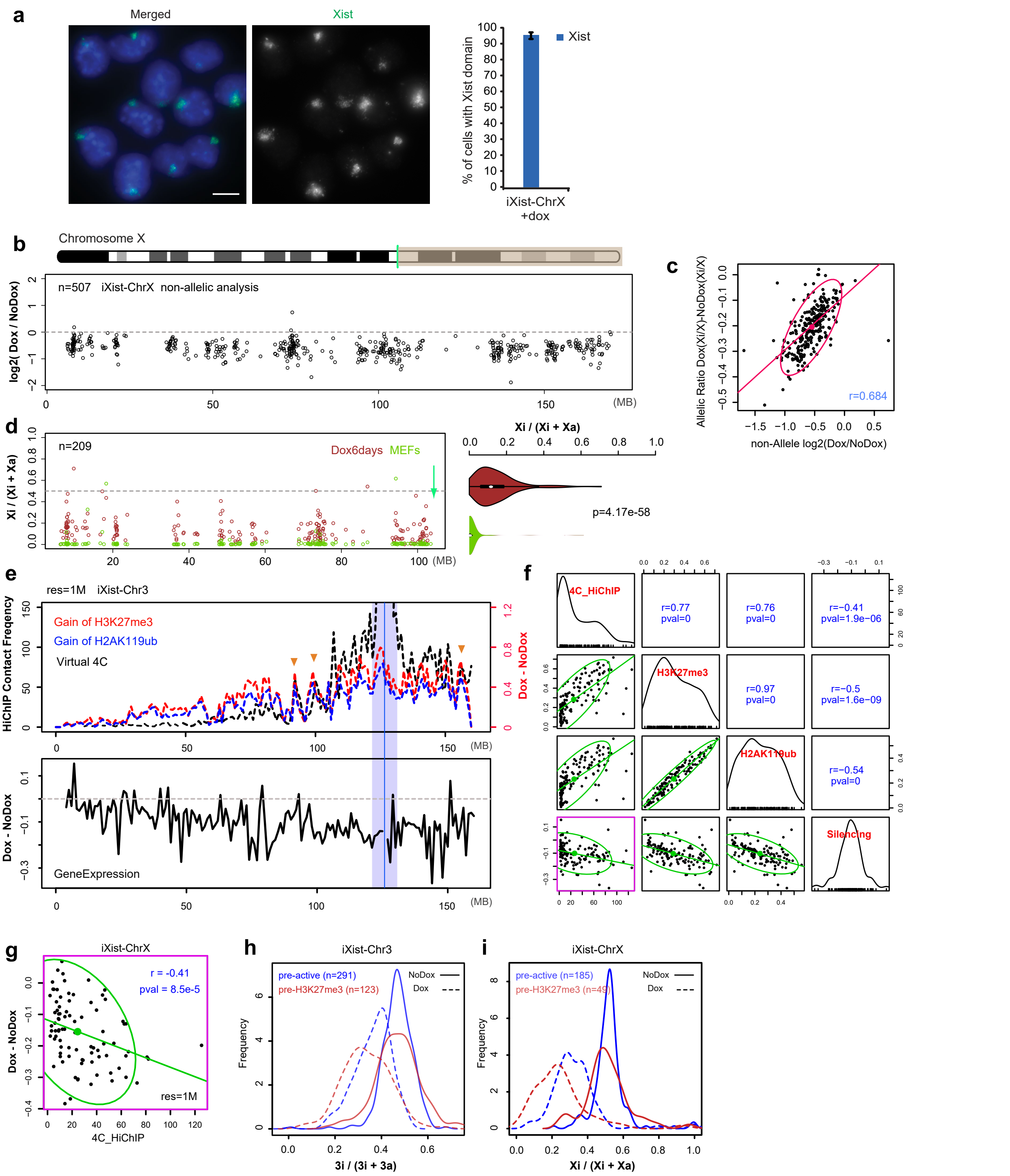


**Supplementary Fig. 1. Supporting data for Fig. 1.** a, Example of RNA FISH analysis of Xist RNA (green) in iXist-ChrX cells after 1 day of doxycycline induction. DNA is counterstained with DAPI (blue). Bar chart shows proportion of cells with Xist RNA domains based on scoring >200 cells in four independent experiments. Error bar indicates standard deviation. b, Gene silencing in iXist-ChrX cells after 1 day of Xist RNA induction as determined by non-allelic analysis**.** Grey shading on ChrX ideogram highlights distal region where informative SNPs are not present. Mean value of allelic ratio for each gene was calculated from biological replicates as detailed in STAR methods. Green line on ChrX ideogram indicates location of the Xist locus. c, Correlation analysis comparing genes proximal to Xist in iXist-ChrX cells analysed either by allelic or non-allelic methodology. Red oval indicates 95% of gene population.

d, Comparison of silencing in iXist-ChrX cells after 6 days of Xist RNA induction with published dataset showing silencing in Cast x FVB XX somatic (MEF) line (Gdula et al., 2018), shown for genes proximal to Xist locus. Mean value of difference of allelic ratio for each gene with an informative SNP was calculated from biological replicates as detailed in STAR methods. Data are summarised in the violin plot to the right. p-values are calculated using one-sided Wilcoxon rank sum test. e, Representation of iXist-Chr3 Xist transgene locus (blue line) 4C-HiChIP contact frequency (black dashed line) together with allelic H3K27me3 and H2AK119ub gain following 1 day of Xist RNA i nduction. Red arrowheads indicate selected highly correlated regions. Allelic silencing is plotted below. f, Correlation analysis showing relationship between 4C-HiChIP, gain of H3K27me3 and H2AK119ub, and allelic silencing in iXist-Chr3 cells after 1 day of Xist RNA induction. Correlation for 4C-HiChIP with silencing (r=-0.41) is highlighted (red box). g, Correlation between allelic silencing and proximity to Xist locus (determined from published 4C-HiChIP data) in iXist-ChrX cells after 1 day of Xist RNA induction. Green oval indicates 95% of gene population. h, i, Allelic repression with/without 1 day of Xist RNA induction determined for Chr3 (H) and ChrX (I) genes were grouped as having pre-existing active (pre-active) or H3K27me3 (pre-H3K27me3) chromatin signature.

### Supplementary Figure 2

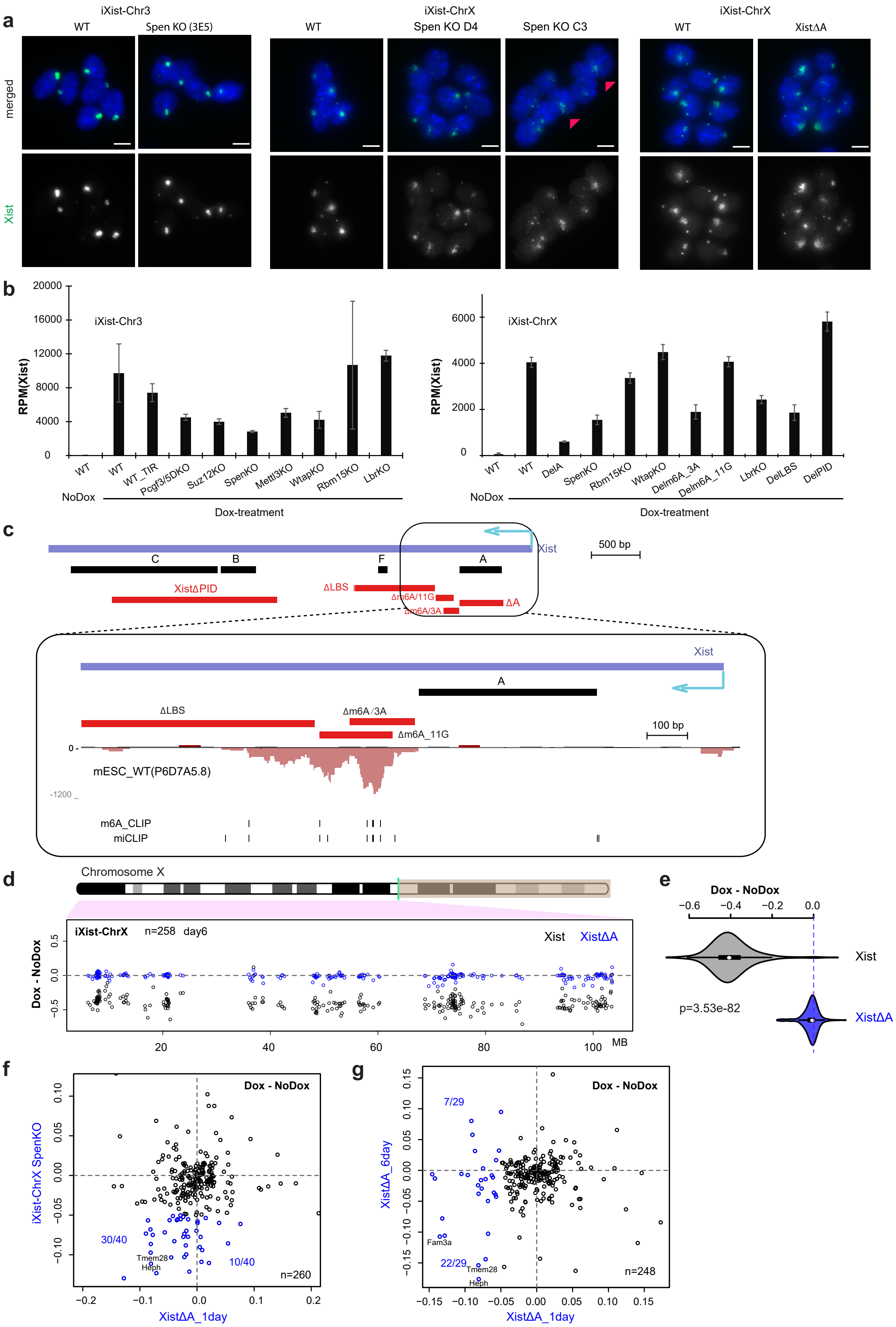

**Supplementary Fig. 2. Supporting data for Fig. 2.** a, Representative images of RNA FISH analysis showing Xist RNA domains (green) in iXist-Chr3, iXist ChrX and derivative mutant cell lines as indicated. DNA is counterstained with DAPI (blue). Scale bar is 10μm. Arrowheads on Spen KO image indicate cells with diffuse Xist RNA domain. b, Xist RNA levels after 1 day of induction determined from ChrRNA-seq experiments and expressed as RPM(Xist) for iXist-Chr3 (left) and iXist-ChrX (right) and derivative mutant mESC lines. Error bars indicate s.e.m. c, Schematic showing the proximal region of the Xist gene indicating the location of different deletions generated in iXist-ChrX cells. The Xist promoter (cyan arrow) is shown for reference. Below zoomed-in region is m6A-seq data indicating the prominent m6A region in Xist RNA. Individual m6A sites determined by m6A-CLiP (Ke et al., 2015), and miCLIP (Linder et al., 2015) are also denoted. d, Allelic silencing across ChrX after 6 days of Xist RNA induction in WT and XistΔA mESCs. Mean value of difference of allelic ratio for each gene with an informative SNP was calculated from biological replicates as detailed in STAR methods. Green line on ChrX ideogram indicates location of the Xist locus. e, Violin plot summarising data in (b). p-values were calculated using one-sided Wilcoxon rank sum test. f, Comparison s howing silencing level for genes that are repressed in Spen KO (blue) after 1 day of Xist RNA induction in XistΔA mESCs. g,. Comparison showing silencing level for genes that are repressed in XistΔA (blue) after 1 day and 6 days of Xist RNA induction in XistΔA mESCs.

#### Supplementary Figure 3

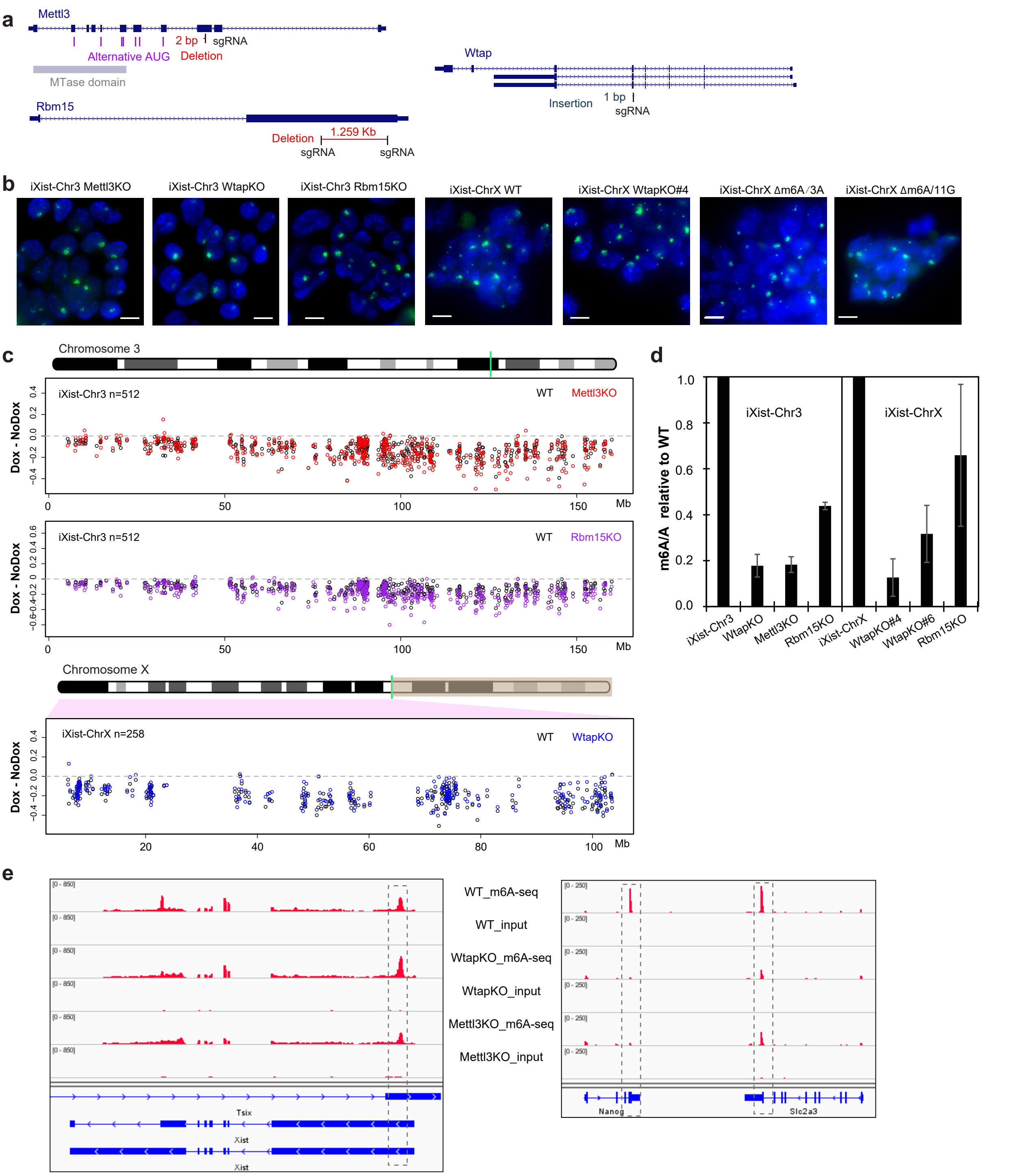


**Supplementary Fig. 3. Supporting data for Fig. 3.** a, Schematics illustrating mutations in Mettl3, Wtap and Rbm15 genes generated by CRISPR/Cas9 mutagenesis. Black vertical bars indicate location of sgRNA complementary sequence. Vertical bars indicate alternative AUG in Mettl3 gene potentially giving rise to truncated catalytically active protein product. b, Representative RNA FISH images showing Xist RNA domains (green) in different iXist-Chr3 and iXist-ChrX mutant lines as indicated. DNA is counterstained with DAPI (blue). Scale bar is 10 μm.

c, Allelic silencing across Chr3 after 1 day of Xist RNA induction in WT compared to Mettl3 mutant (KO) (top), in WT compared to Rbm15 null (KO) iXist-Chr3 mESCs (middle), and for WtapKO across ChrX in iXist-ChrX cells (bottom). Mean value of difference of allelic ratio for each gene with an informative SNP was calculated from biological replicates as detailed in STAR methods. Green lines on Chr3 and ChrX ideograms indicate location of the Xist transgene/locus. d, Levels of m6A in mRNA from iXist-Chr3 and iXist-ChrX mESCs and indicated mutant cell lines determined by LC-MS/MS. Two independent iXist-ChrX WtapKO cell l ines were analysed. Error bars indicate variation between two biological replicates. e, IGV browser screenshots illustrating m6A-seq data across the Xist locus (left) and Nanog and Slc2a3 loci (right) in WT iXist-Chr3 mESCs and derivative Wtap KO and Mettl3 KO as indicated. Dotted boxes indicate major m6A peaks.

**Supplementary Figure 4**

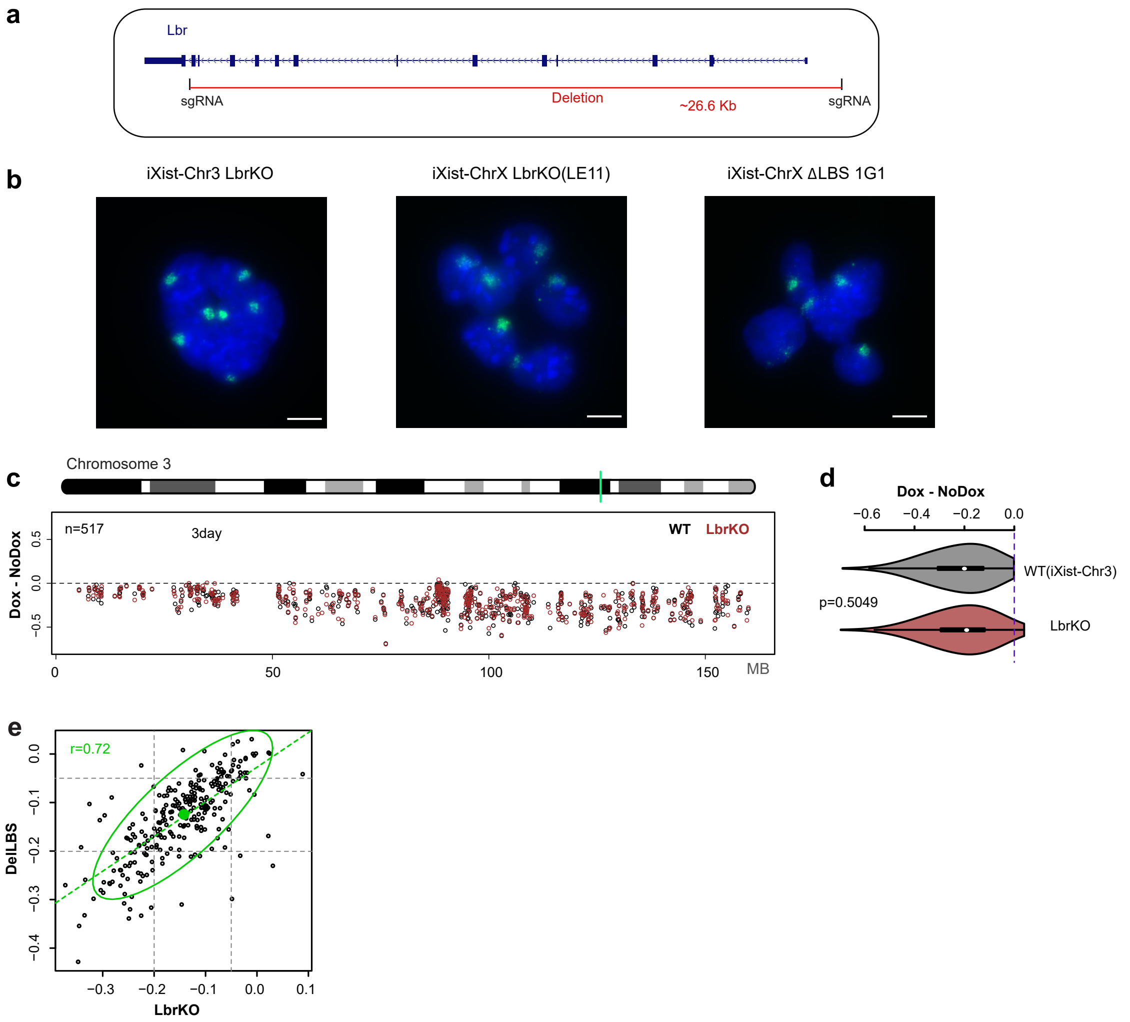


**Supplementary Fig. 4. Supporting data for Fig. 4.** a, Locus map of the Lbr gene showing the location of the deletion generated by CRISPR/Cas9 mutagenesis. b, Representative RNA FISH images showing Xist RNA domains (green) in different iXist-Chr3 and iXist-ChrX mutant lines as indicated. DNA is counterstained with DAPI (blue). Scale bar is 10 μm. c, Allelic silencing across Chr3 after 3 days of Xist RNA induction in WT compared to Lbr null (KO) iXist-Chr3 mESCs. Mean value of difference of allelic ratio for each gene with an informative SNP was calculated from biological replicates as detailed in STAR methods. Green line on Chr3 ideogram indicates location of the Xist transgene. d, Violin plot summarising data from (c). p-values were calculated using one-sided Wilcoxon rank sum test. e, Correlation of gene silencing in Lbr null and XistΔLBS mESCs. Green oval represents 95% of the gene population.

#### Supplementary Figure 5

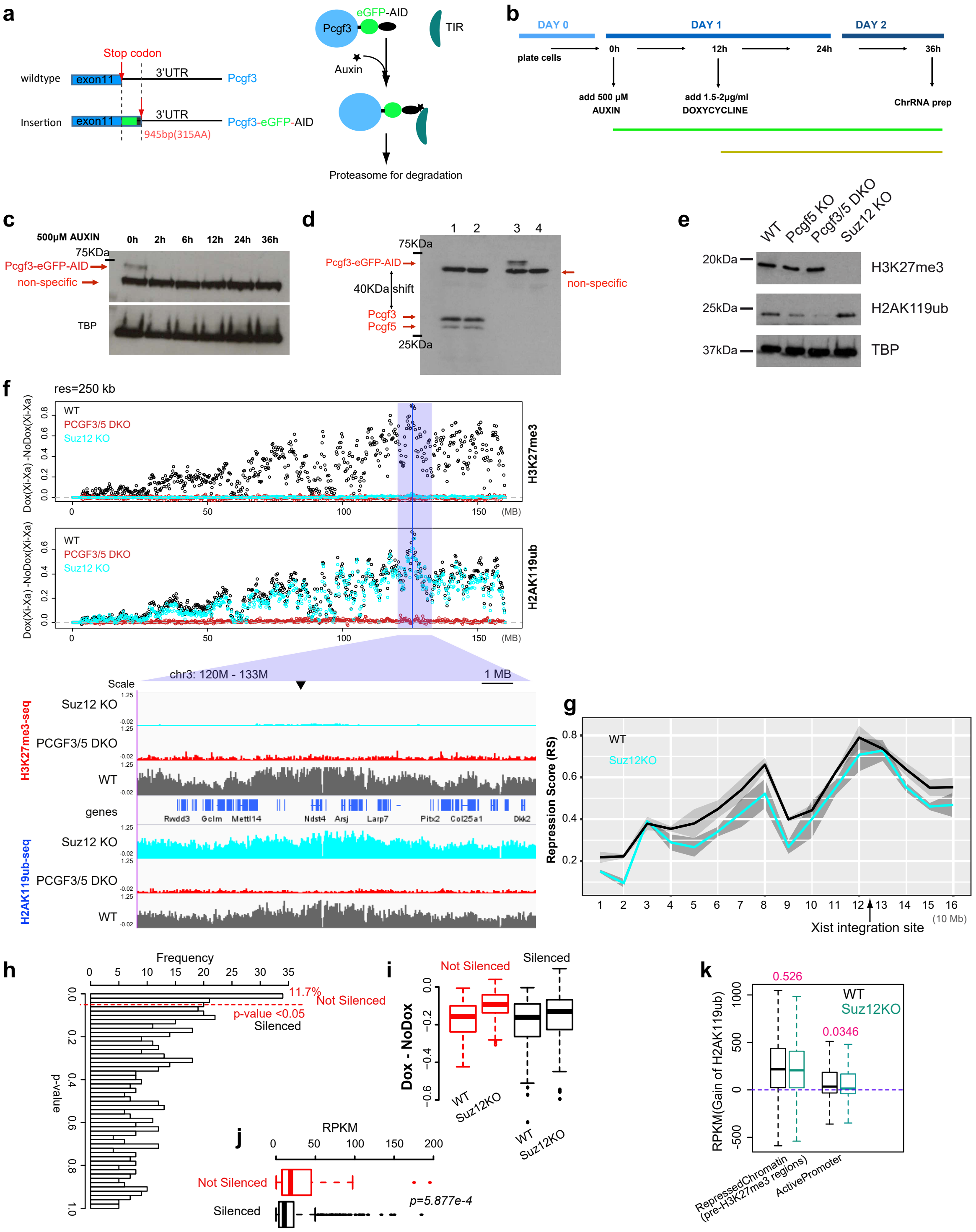


**Supplementary Fig. 5. Supporting data for Fig. 5.** a, Strategy for tagging Pcgf3 exon 11 with an in frame GFP-AID tag. b, Schematic for PCGF3 degradation showing Auxin treatment (green line) and Xist RNA induction (yellow line) time windows. c, Western blot illustrating that degradation of PCGF3 in Pcgf3/5 DKO mESCs occurs within 2 hours of addition of Auxin. TBP is a loading control. d, PCGF3 + PCGF5 western blot in WT iXist-Chr3 mESCs (lanes 1 and 2), and in Pcgf3/5 DKO mESCs (lanes 3 and 4). Auxin was added for 36h (lanes 2 and 4). e, Western blot showing global H3K27me3 and H2AK119ub levels in cell lines as indicated. DKO is post-Auxin treatment. TBP is a loading control.

f, Gain of H3K27me3 (top) and H2AK119ub (bottom) on Chr3 determined by calibrated allelic ChIP-seq after 1 day of Xist RNA induction in WT, Pcgf3/5 DKO and Suz12 KO iXist-Chr3 mESCs as indicated. The location of the Xist transgene is shown with a blue vertical line. IGV browser screenshot zoomed to 13Mb around the Xist transgene is shown below. g, Allelic silencing illustrated as repression score (Pintacuda et al, 2017), for WT and Suz12 KO mESCs after 3 day of Xist RNA induction. h, Frequency of genes that are not silenced ($p<0.05$) in Suz12 KO compared to WT mESCs after 3 days of Xist RNA induction. i, Boxplot illustrating allelic silencing in Suz12 KO for genes classified in (H) as Not silenced and Silenced. j, Boxplot illustrating Not Silenced and Silenced genes in Suz12 KO plotted against expression level in mESCs. p-value is calculated from two-sided Wilcoxon rank sum test. k, Boxplot illustrating gain of H2AK119ub at genes with pre-existing H3K27me3 (pre-H3K27me3) or active (pre-active) chromatin marks in WT and Suz12 KO mESCs after 1 day of Xist RNA induction. p-value is calculated from two-sided Wilcoxon rank sum test.

#### Supplementary Figure 6

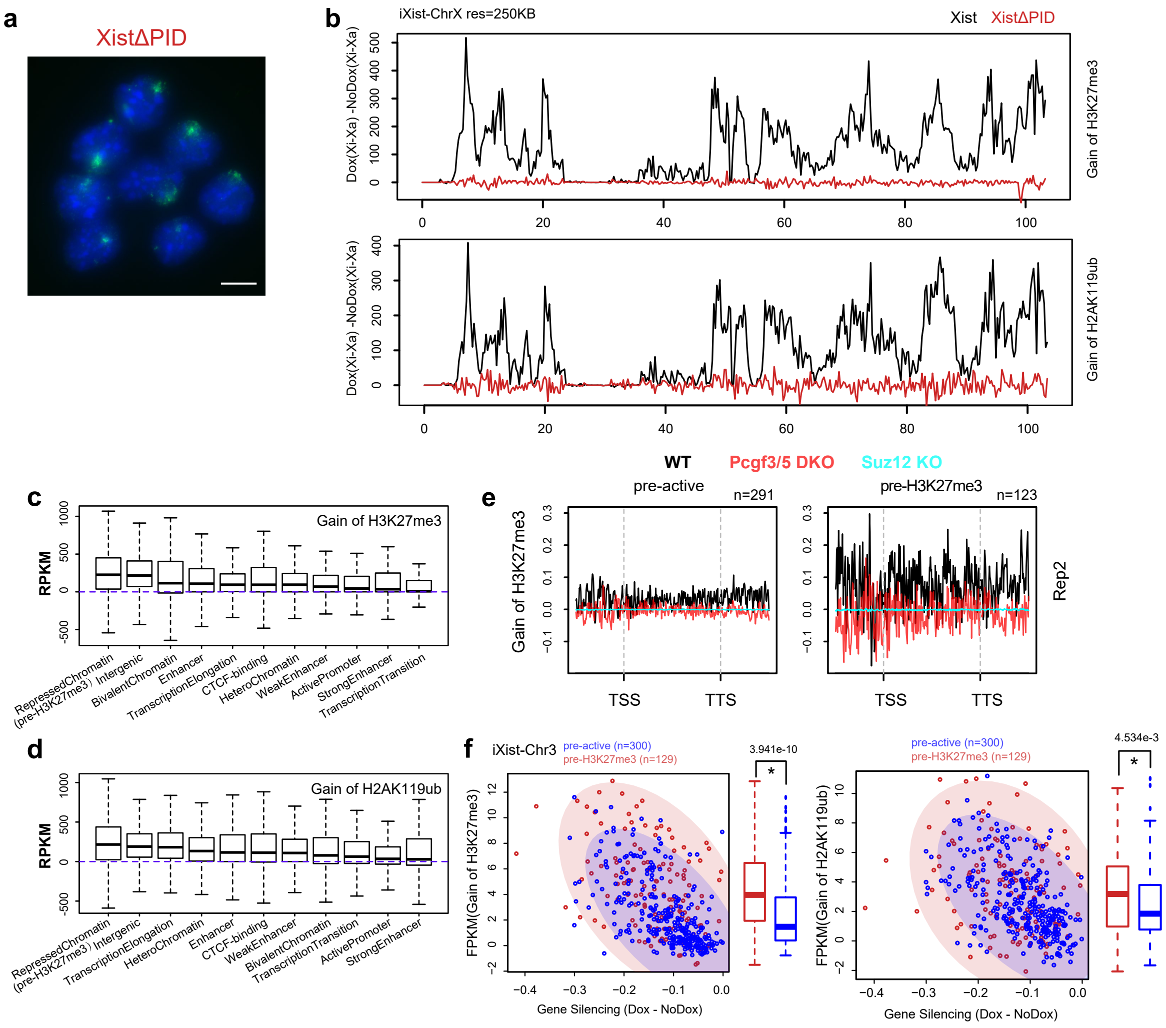


**Supplementary Fig. 6. Supporting data for Fig. 5 and 6.** a, Representative image of RNA FISH analysis showing Xist RNA domains (green) in iXist-ChrX XistΔPID mESCs. DNA is counterstained with DAPI (blue). Scale bar is 10µm. b, Gain of H3K27me3 (top) and H2AK119ub (bottom) in 500kb bins on Xist proximal ChrX determined by calibrated allelic ChIP-seq after 1 day of Xist RNA induction in iXist-ChrX and XistΔPID mESCs. c, d, ChromHMM analysis showing gain of H3K27me3 (C) and H2AK119ub (D) at different regions defined by epigenetic chromatin state in iXist-Chr3 mESCs after 1 day of Xist RNA induction. e, Metaprofile complementing Fig. 6g, h using a different biological replicate. f, Scatterplots and boxplots illustrating increased gain of H3K27me3 (left) and H2AK119ub (right) in gene promoters with pre-existing H3K27me3 (pre-H3K27me3) compared with active (pre-active) chromatin marks in iXist-Chr3 mESCs after 1 day of Xist RNA induction.

#### Supplementary Figure 7

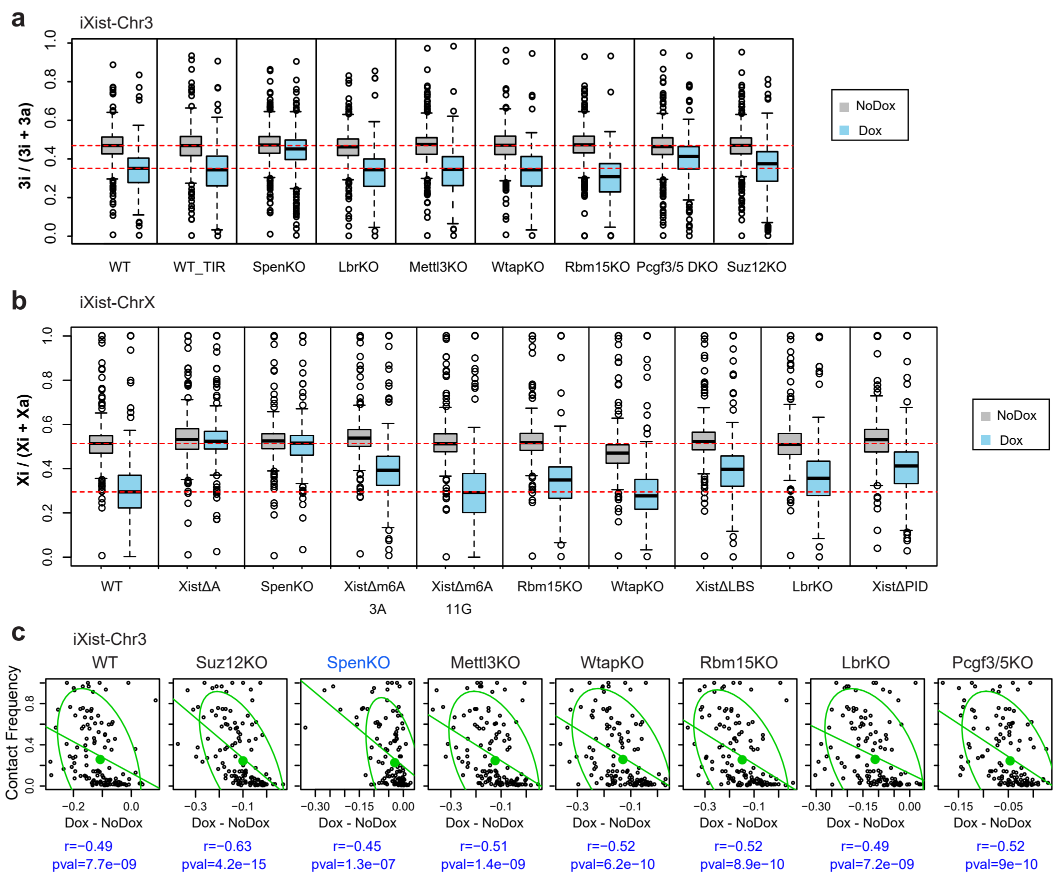


**Supplementary Fig. 7. Supporting data for Fig. 7.** a, Boxplots summarising allelic silencing after 1 day of Xist RNA induction in iXist-Chr3 and different mutant derivative mESCs. WT values represent an average of all determinations except for Pcgf3/5 DKO where the WT-TIR cell line is the appropriate control. b, As in (a) for iXist-ChrX.
c, Correlation analysis for topological distance and silencing efficiency for WT and selected mutant mESCs in iXist-Chr3 model. 95% of the population is encircled. Spearman correlation and p-values are indicated below.

**Supplementary Table 1. Cell Lines used in this study and validation methods used to characterized each line**

| | |
|---|---|
| mESC: iXist-Chr3, XY | WB, RNA-FISH, RNAseq |
| mESC: iXist-Chr3 LBR KO, XY | gPCR, SS, SB, RNAseq |
| mESC: iXist-Chr3 METTL3 KO, XY | gPCR, SS, WB, RNA-FISH, RNAseq |
| mESC: iXist-Chr3 PCGF3-AID PCGF5 KO, XY | gPCR, SS, WB |
| mESC: iXist-Chr3 RBM15 KO, XY | SS, WB, RNAseq |
| mESC: iXist-Chr3 SPEN KO, XY | gPCR, SS, SB, RNAseq |
| mESC: iXist-Chr3 WTAP KO, XY | gPCR, SS, WB, IF, RNA-FISH, RNAseq |
| mESC: iXist-ChrX, XX | gPCR, SS, SB, IF, RNA-FISH, mFISH, RNAseq, ChIPseq |
| mESC: iXist-ChrX LBR KO, XX | gPCR, SS, SB, RNAseq |
| mESC: iXist-ChrX RBM15 KO, XX | gPCR, WB, RNAseq |
| mESC: iXist-ChrX SPEN KO, XX | gPCR, SS, SB, RNAseq |
| mESC: iXist-ChrX WTAP KO B2.4, XX | gPCR, SS, IF, RNA-FISH, RNAseq |
| mESC: iXist-ChrX WTAP KO B2.6, XX | gPCR, SS, IF, RNA-FISH, RNAseq |
| mESC: iXist-ChrX Xist ΔA, XX | gPCR, SS, RNAseq |
| mESC: iXist-ChrX XistΔLBS KO, XX | gPCR, SS, SB, RNAseq |
| mESC: iXist-ChrX XistΔm6A 3A, XX | gPCR, RD, SS, RNA-FISH, RNAseq |
| mESC: iXist-ChrX Xist Δm6A 11G.7, XX | gPCR, RD, SS, RNA-FISH, RNAseq |
| mESC: iXist-ChrX Xist ΔPID, XX | gPCR, SS, RNAseq |

**Supplementary Table 2. List of Oligonucleotides**

| Oligonucleotide | Source | Used for |
|---|---|---|
| LBR_gRNA1, ATGATGTTGAATTCGTATAC | This study | CRISPR KO |
| LBR_gRNA2, GCCTACCTTATGAGCGGGTT | This study | CRISPR KO |
| LBR_F, GTTAAATGTTAAGGAAAAGCAGG | This study | gPCR, sequencing |
| LBR_R, TCACCCACTGAAAGCCAAGC | This study | gPCR, sequencing |
| METTL3_gRNA, ACGCCGTTTCTGCCCTGCGATGG | This study | CRISPR KO |
| METTL3_F, CAGCGTGCTTCCTTGACTTCTT | This study | gPCR, sequencing |
| METTL3_R, TCCAAGATGATGCACATCCTACTCT | This study | gPCR, sequencing |
| PCGF3_gRNA1, CCAGCACGCGTTTACAGAGGA | This study | CRISPR HR |
| PCGF3_F, GAAAGGTTGAACGTGCACCT | This study | gPCR, sequencing |
| PCGF3_R, GCGAGCTGTTTCCTCAAGTG | This study | gPCR, sequencing |
| PCGF5_gRNA1, GCTCCTCTGCTTCAACTGCT | This study | CRISPR KO |
| PCGF5_gRNA2, GATCAAGCCCACGACAGTGACGG | This study | CRISPR KO |
| PCGF5_F, TGTTTACAGAGAGGAAGCGCC | This study | gPCR, sequencing |
| PCGF5_R, TGGCCTTGGTACACATATAGC | This study | gPCR, sequencing |
| RBM15_gRNA1, TCACCGCCACGGGAGCGCTT | This study | CRISPR KO |
| RBM15_gRNA2, GCACGAGAATTTGACCGATT | This study | CRISPR KO |
| RBM15_F, GGAGTCCAAGATGGCGGCGTG | This study | gPCR, sequencing |
| RBM15_R, CACTAGTTCATAGTGGGTCAAGG | This study | gPCR, sequencing |
| SPEN_gRNA3, GGGGTGTCTCCTGCGCATT | Monfort et al, 2015 | CRISPR KO |
| SPEN_gRNA5, CGGACAAGACATTACGATC | Monfort et al, 2015 | CRISPR KO |
| SPEN_F, CAGAAAGAGGCGAGGCGTAAAG | This study | gPCR, sequencing |
| SPEN_R, GCGCTCCAGCCGAGCCTTCTC | This study | gPCR, sequencing |

| | | |
|---|---|---|
| TIGRE_gRNA, ACTGCCATAACACCTAACTT | M.Houlard | CRISPR HR |
| TIGRE_rtTA_F, AAGGGGGAGGATTGGGAAGAC | This study | gPCR, sequencing |
| TIGRE_R, GTGATCCACGGTGATCCACA | M.Houlard | gPCR, sequencing |
| WTAP_gRNA, TACTCAGCAAACGATGTGACTGG | This study | CRISPR KO |
| WTAP_F, GTAGTCCCTTGATGCAAAGTA | This study | gPCR, sequencing |
| WTAP_R, AACAGTTTTGTTTTGAGGAG | This study | gPCR, sequencing |
| Xist_TRE_gRNA1, TAACTGATCCGCGGCGCTGA | This study | CRISPR HR |
| Xist_TRE_gRNA2, GCACGCCTTTAACTGATCCG | This study | CRISPR HR |
| Xist ΔA_gRNA, GATCAGTTAAAGGCGTGCAA | This study | CRISPR HR |
| Xist ΔA_F, TACCCGGGGATCCTCTAGTC | This study | gPCR, sequencing |
| Xist ΔA_R, CGGTGTCCTAATTCTTGGCG | This study | gPCR, sequencing |
| XistΔLBS_gRNA1, TTTAGCTAGCGCAGCGCAAT | This study | CRISPR HR |
| XistΔLBS_gRNA2, AAGCATGCGCTCTCCCGACC | This study | CRISPR HR on 129 allele |
| XistΔLBS_F, ACGGCTATTCTCGAGCCAGTT | This study | gPCR, sequencing |
| XistΔLBS_R, GGACTCCAAAGTAACAATTC | This study | gPCR, sequencing |
| XistΔm6A_gRNA1, TTTTTTCGAAGTGCCTGCCCAGG | This study | CRISPR HR on 129 allele |
| XistΔm6A_F, TTTTTTTTCACGGCCCAACGGGGCG | This study | gPCR, sequencing |
| XistΔm6A_R, ATACCGCACCAAGAACTTGAGCC | This study | gPCR, sequencing |
| Xist ΔPID_gRNA, TTGAGGTTCTATACCAGTTC | This study | CRISPR HR |
| Xist ΔPID_F, TCAAGCGGTTCTCTAAGCCT | This study | gPCR, sequencing |
| Xist ΔPID_R, : GATGATACCCTCCCATGGCA | This study | gPCR, sequencing |

**Supplementary Table 3. List of Antibodies used in the study**

| Antibody | Company | Cat. number |
|---|---|---|
| WTAP, Rabbit, WB, IF | Proteintech | Cat#10200-1-AP |
| METTL3, Rabbit, WB, IF | Abcam | Cat#ab195352 |
| m6A, Rabbit, M6A -seq | Synaptic systems | Cat#202003 |
| H2AK119ub, Rabbit, ChIP | Cell Signalling | Cat#8240S |
| H3K27me3, Rabbit, ChIP | Diagenode | Cat# C15410069 |
| H3K27me3, Mouse, IF | Active Motif | Cat# 61017 |
| RBM15, Rabbit, WB | Proteintech | Cat# 10587-1-AP |
| PCGF3+PCGF5, WB | Abcam | Cat#ab201510 |
| Alexa 568 anti-mouse IgG, Goat, IF | Life Technologies | cat#A11031 |
| Alexa 488 anti-mouse IgG , IF | Life Technologies | cat#A11029 |
| Alexa 568 anti-rabbit IgG, Goat, IF | Life Technologies | cat#A11034 |
| Alexa 488 anti-rabbit IgG, Goat, IF | Life Technologies | cat#A11008 |
| Anti-rabbit Ig, HPR, Donkey, WB | Amersham | cat#NA934V |
| Anti-mouse IgG, HRP, Sheep, WB | Amersham | cat#NXA931 |

**Supplementary Table 4. Biological Replicate RNA Sequencing**

| Cell lines | ES/DIF, treatment time | +Dox Replicates | No Dox Replicates | Sequencing |
|---|---|---|---|---|
| iXist-Chr3 | ES, 1 day | 3 | 4 | ChrRNA |
| iXist-Chr3_Pcgf5KO_TIR | ES, 1 day | 2 | 1 | ChrRNA |
| iXist-Chr3_Pcgf5KO_TIR_B7 | ES, 1 day | 2 | 1 | ChrRNA |
| iXist-Chr3_Pcgf5KO_TIR_C10 | ES, 1 day | 2 | 1 | ChrRNA |
| iXist-Chr3_Suz12KO | ES, 1 day | 2 | 1 | ChrRNA |
| iXist-Chr3_SpenKO#2E5 | ES, 1 day | 1 | 1 | ChrRNA |
| iXist-Chr3_SpenKO#2G9 | ES, 1 day | 1 | 1 | ChrRNA |
| iXist-Chr3_SpenKO#3E5 | ES, 1 day | 1 | 1 | ChrRNA |
| iXist-Chr3_SpenKO#3H6 | ES, 1 day | 1 | 1 | ChrRNA |
| iXist-Chr3_LbrKO#4C7A | ES, 1 day | 1 | 1 | ChrRNA |
| iXist-Chr3_LbrKO#4C8E | ES, 1 day | 1 | 1 | ChrRNA |
| iXist-Chr3_LbrKO#4F9F | ES, 1 day | 1 | 1 | ChrRNA |
| iXist-Chr3_Mettl3KO | ES, 1 day | 2 | 1 | ChrRNA |
| iXist-Chr3_WtapKO | ES, 1 day | 2 | 1 | ChrRNA |
| iXist-Chr3_Rbm15KO | ES, 1 day | 2 | 1 | ChrRNA |
| iXist-Chr3 | DIF, 3 days | 3 | 3 | ChrRNA |
| iXist-Chr3_Suz12KO | DIF, 3 days | 3 | 3 | ChrRNA |
| iXist-Chr3_LbrKO#4C7A | DIF, 3 days | 1 | 1 | ChrRNA |
| iXist-Chr3_LbrKO#4C8E | DIF, 3 days | 1 | 1 | ChrRNA |
| iXist-Chr3_LbrKO#4F9F | DIF, 3 days | 1 | 1 | ChrRNA |
| iXist-ChrX | ES, 1 day | 6 | 5 | ChrRNA |
| iXist-ChrX_XistΔA | ES, 1 day | 2 | 1 | ChrRNA |
| iXist-ChrX_XistΔPID | ES, 1 day | 2 | 1 | ChrRNA |
| iXist-ChrX_XistΔLBS#1G1 | ES, 1 day | 1 | 1 | ChrRNA |
| iXist-ChrX_XistΔLBS#1C7 | ES, 1 day | 1 | 1 | ChrRNA |
| iXist-ChrX_LbrKO#LE11 | ES, 1 day | 1 | 1 | ChrRNA |
| iXist-ChrX_LbrKO#LH4 | ES, 1 day | 1 | 1 | ChrRNA |
| iXist-ChrX_Rbm15KO#E10 | ES, 1 day | 3 | 3 | ChrRNA |
| iXist-ChrX_Rbm15KO#A8 | ES, 1 day | 4 | 3 | ChrRNA |
| iXist-ChX_SpenKO#C3 | ES, 1 day | 1 | 1 | ChrRNA |
| iXist-ChX_SpenKO#C4 | ES, 1 day | 1 | 1 | ChrRNA |
| iXist-ChX_SpenKO#D4 | ES, 1 day | 1 | 1 | ChrRNA |
| iXist-ChrX_Δm6A/3A | ES, 1 day | 2 | 2 | ChrRNA |
| iXist-ChrX_Δm6A/11G | ES, 1 day | 2 | 2 | ChrRNA |
| iXist-ChrX_WtapKO#4 | ES, 1 day | 2 | 1 | ChrRNA |
| iXist-ChrX_WtapKO#6 | ES, 1 day | 2 | 1 | ChrRNA |
| iXist-ChrX | ES, 6 days | 1 | 1 | ChrRNA |
| iXist-ChrX_XistΔA | ES, 6 days | 1 | 1 | ChrRNA |
| iXist-ChrX_XistΔPID | ES, 6 days | 1 | 1 | ChrRNA |

| | | | | |
|---|---|---|---|---|
| iXist-ChrX | DIF, 6 days | 3 | 3 | ChrRNA |
| iXist-ChrX_XistΔA | DIF, 6 days | 3 | 3 | ChrRNA |
| iXist-ChrX_XistΔPID | DIF, 6 days | 3 | 3 | ChrRNA |
